# Biochemical mechanisms driving rapid fluxes in C_4_ photosynthesis

**DOI:** 10.1101/387431

**Authors:** A Bräutigam, U Schlüter, MR Lundgren, S Flachbart, O Ebenhöh, G Schönknecht, PA Christin, S Bleuler, JM Droz, CP Osborne, APM Weber, U Gowik

## Biochemical mechanisms driving rapid fluxes in C_4_ photosynthesis

Sufficient flux through pathways is critical to their function within the metabolic network. A high flux cycle called C_4_ photosynthesis evolved in some plants to combat the slow speed and low specificity of the plant’s carbon fixation enzyme Rubisco^1^. In these plants, flux through the C_4_ photosynthetic cycle eclipses flux through Rubisco making it the highest flux *in planta*^2,3^. Engineering efforts are underway to transfer this high flux pathway to crops using the ancestral C_3_ photosynthetic type, such as rice, to improve yield and increase water and nitrogen use efficiencies^4^. Most of the enzymatic and transport steps of the C_4_ cycle have been characterized^5^. However, well-established features, including altered chloroplast structure, diversity in transfer acids, and the underlying mechanisms driving high flux through this biochemical CO_2_ pump^2,3^ remain unexplained.

We identify three drivers of high fluxes through alternative decarboxylation subsystems in C_4_ plants: For NAD^+^-and NADP^+^-based malic enzymes, kinetic modeling demonstrates that the oxidation state of mitochondria and chloroplasts, respectively, drive rapid decarboxylation. For phospho*enol*pyruvate carboxykinase, RNA-seq and enzyme activity measurements suggest the presence of a new C_4_ cycle enzyme driving fast decarboxylation. These observations not only suggest a hitherto unknown C_4_ cycle but also provide mechanistic explanations for decades-old anatomical^6,7^ and biochemical observations^8^.

C_4_ photosynthesis has evolved to overcome kinetic constraints of the carbon fixation enzyme Rubisco^1^ (Ribulose-1,5-bisphosphate carboxylase oxygenase) by enriching CO_2_ at its active site^8^. Rubisco is the carboxylation enzyme common to all plants and, under current atmospheric conditions, it catalyzes both a productive carboxylation reaction and a wasteful oxygenation reaction^1,9^. Enriching CO_2_ suppresses the wasteful side reaction at the cost of synthesizing and transferring metabolite in a cyclic biochemical CO_2_ pump. Most of the enzymatic and transport steps of this C_4_ cycle have been characterized^5,8^. C_4_ species evolved increased transcript and protein abundance of C_4_ cycle enzymes and transporters to support a high flux pathway^10–13^.

C_4_ cycles with different decarboxylation enzymes operate using the same principles. The cycle pre-fixes CO_2_ in mesophyll cells to a three-carbon acceptor, PEP, resulting in a four-carbon organic acid, which is modified and transferred to the site of Rubisco in bundle sheath cells^14^ (Figure 1). Decarboxylation then liberates CO_2_ and this reaction enriches CO_2_ at the active site of Rubisco, if decarboxylation exceeds the Rubisco reaction by 20% to accommodate leakage^2,3^. The resulting C_3_ acid is transferred back to the mesophyll site of pre-fixation to regenerate the C_3_ acceptor (Figure 1). Regeneration of PEP requires the hydrolysis of two ATP molecules and drives the carboxylation side of the cycle^8^. Transfer of the C_4_ pre-fixation product from the mesophyll to bundle sheath cells, and return of the C_3_ decarboxylation product occurs by diffusion driven by concentration gradients^15,16^. This requires low C_4_ acid concentration and high C_3_ acid concentration in the bundle sheath cells to sustain the necessary transport rates (Figure 1). These constraints imply that accelerating decarboxylation by increased substrate (C_4_ acid) or decreased product (C_3_ acid) concentration in the bundle sheath is not a feasible option. Moreover, since the cycle functions to increase CO_2_ concentration, the decarboxylation velocity also cannot be increased via lowering the CO_2_ concentration. These constraints raise major uncertainties about how a rapid flux of CO_2_ to Rubisco is generated and maintained in C_4_ photosynthesis. We hypothesized that anatomical and/or biochemical adaptations modify the bundle sheath environment to drive the decarboxylation rate towards maximal velocity despite these unfavorable constraints. To identify the adaptations, we modeled decarboxylation with kinetic and conceptual models.

**Figure 1:**
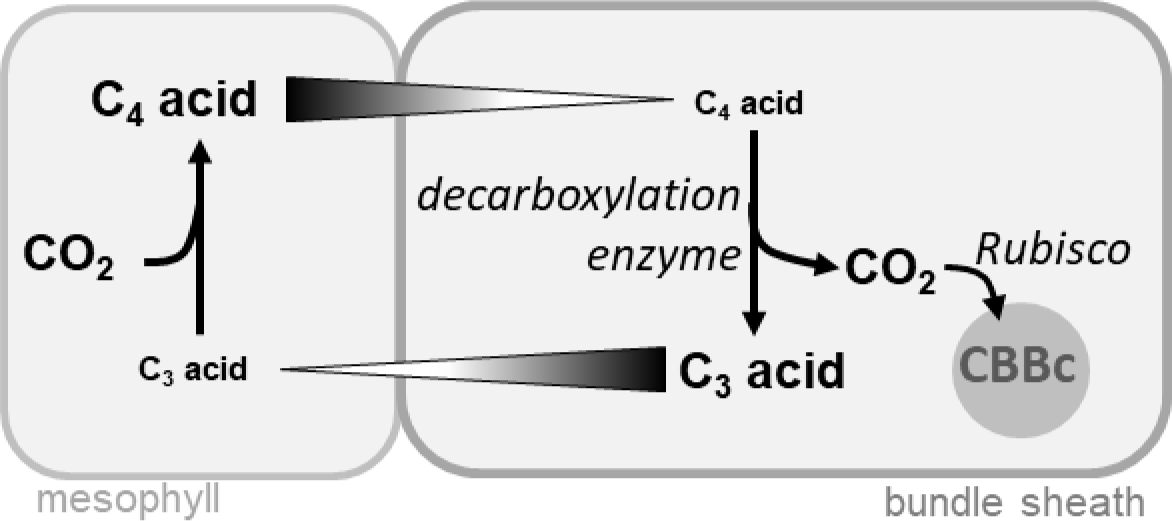
**The schematic representation of a generalized C_4_ cycle** shows the necessary concentration gradients in the C_4_ cycle and the resulting constraints for decarboxylation; flux through the decarboxylation enzyme needs to exceed flux through Rubisco to achieve CO_2_ enrichment.

### The NADP-ME subtype

In this subtype, NADP-dependent malic enzyme (NADP-ME) decarboxylates malate to pyruvate and CO_2_ while reducing NADP^+^ in chloroplasts; malate is the main intermediate transfer acid^8,18^. The constraints on decarboxylation via NADP-ME were analyzed with a kinetic model. This model was derived from the general kinetic reaction for the random binding of ligands^19^, after conversion to a form that included the equilibrium constant K_eq_ for reversible reactions^3^. The kinetic model was parametrized for *Zea mays* NADP-ME using metabolite concentrations from currently available data from the literature. The velocity of decarboxylation was calculated as a function of the NADP^+^/NADPH ratio and compared to the known photosynthetic rate of mature *Z. mays* leaves (Figure 2A). When the concentration of NADP^+^ is very small, the reaction occurs in the carboxylation direction. With greater NADP^+^/NADPH ratio, the decarboxylation rate increases (Figure 2A, Supplemental R Code) indicating that oxidized redox poise in the chloroplast stroma favors high velocity of decarboxylation. Bundle sheath chloroplasts indeed achieve this more oxidized state via two mechanisms: First, the NADPH generated by NADP-ME based decarboxylation is consumed by the Calvin Benson Bassham cycle (Figure 2B) and secondly, the production of NADPH through the photosynthetic electron transfer chain is limited by abolishing or reducing photosystem II activity^20^ in bundle sheath cells (Figure 2B). This second mechanism results in the anatomically differentiated chloroplasts common to C_4_ species deploying NADP-ME as a decarboxylase^7,21,22^ which represent a derived state compared to chloroplasts in C_3_ plants. Our model mechanistically explains limited photosystem II activity as a specific adaptation to drive high velocity decarboxylation by NADP-ME in the chloroplasts which operates by changing the redox poise (Figure 2B).

**Figure 2:**
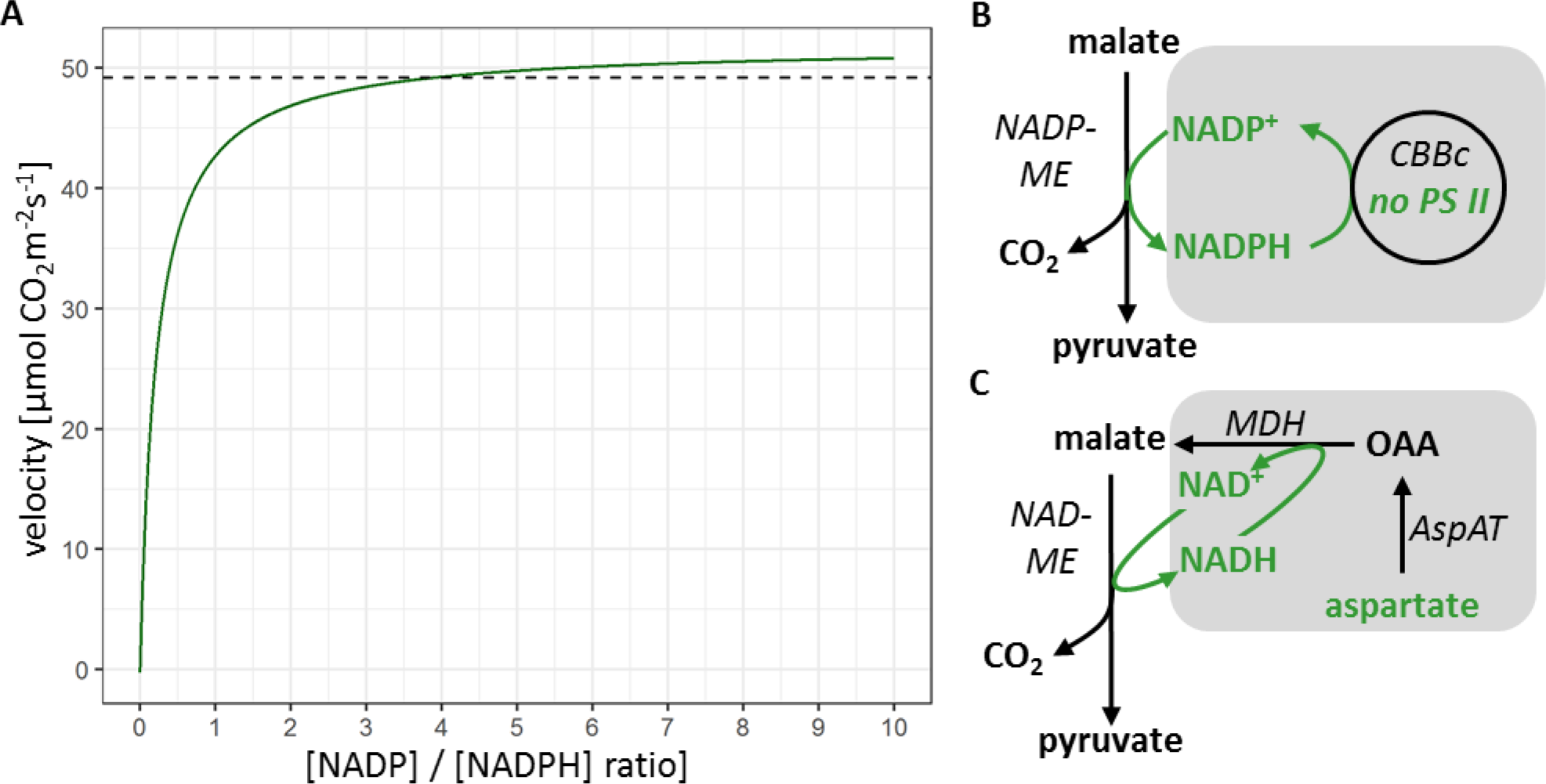
**Decarboxylation model for malic enzyme; (A)** velocity of NADP-ME-based carboxylation depends on the [NADP^+^]/[NADPH] ratio at the site of the enzyme as calculated by the model parametrized for **Zea mays**NADP-ME; the necessary velocity is represented as a dashed line and was calculated from carbon fixation rate time overcycling requirement^17^; **(B)** NADP-ME based decarboxylation in the chloroplast is driven by NADPH consumption of the Calvin Benson Bassham cycle (CBBc) and the absence of photosystem II (PS II), driver represented as grey box; **(C)** NAD-ME based decarboxylation in the mitochondria is driven by provision of NAD through coupling to the oxaloacetate (OAA) to malate reaction, driver represented as grey box;

### The NAD-ME subtype

In this subtype, NAD-dependent malic enzyme (NAD-ME) decarboxylates malate to pyruvate and CO2 while reducing NAD^+^ in mitochondria; aspartate is the main intermediate transfer acid^8,18^. The model for NAD-ME based decarboxylation is conceptual and derived from the NADP-ME model, since neither metabolite concentrations nor enzyme parameters are available. One may qualitatively derive from the model parametrized for NADP-ME (Figure 2A) that a higher NAD^+^/NADH ratio in the mitochondria will drive a faster decarboxylation reaction in NAD-ME plants since standard redox potentials of the pairs NAD^+^/NADH and NADP^+^/NADPH are similar. C_3_ mitochondria display redox poise favorable to decarboxylation with NAD^+^/NADH ratios between three and ten^23^. Bundle sheath mitochondria maintain this favorable redox poise: although NAD-ME-based decarboxylation of malate in C_4_ mitochondria yields NADH, it is regenerated to NAD^+^ via the reduction of oxaloacetate within the C_4_ cycle (Figure 2C) if aspartate is the transfer acid. The reduction of OAA derived from aspartate is stoichiometrically coupled to decarboxylation resulting in maintenance of redox poise. It was previously unclear, why NAD-ME plants transfer CO_2_ via aspartate and not malate^8^. Our model explains the preference as a specific adaptation to regenerate NAD^+^ and drive high velocity decarboxylation by NAD-ME via the redox poise of mitochondria (Figure 2C). In principle, using aspartate as the transfer acid should also contribute favorably to redox poise in NADP-ME plants. Indeed, in the C_4_ eudicot *Flaveria bidentis* PS II function is reduced but not absent^24^ and considerable aspartate flux is detectable^25^.

### The PEP-CK subtype

In this subtype, phosphoenolpyruvate carboxykinase (PEP-CK) decarboxylates oxaloacetate to phosphoenolpyruvate (PEP) and CO_2_ while consuming ATP in the cytosol. In current models PEP-CK activity is accompanied by substantial NAD- or NADP-ME activity indicating that they use mixed decarboxylation systems ^8,26–28^. We analyzed a species that relies predominantly on PEP-CK activity, *Alloteropsis semialata* subs. *semialata*^29–31^ that – according to the current models^8,26,28^ – should not exist in nature. Relying solely, or even predominantly, on PEP-CK is theoretically impossible^26,28^, since the absence of malic enzyme results in imbalanced amino- and phosphate groups^26,28^ as well as in an incomplete C_4_ cycle with an unresolved gap between PEP and pyruvate (Figure 3A).

**Figure 3:**
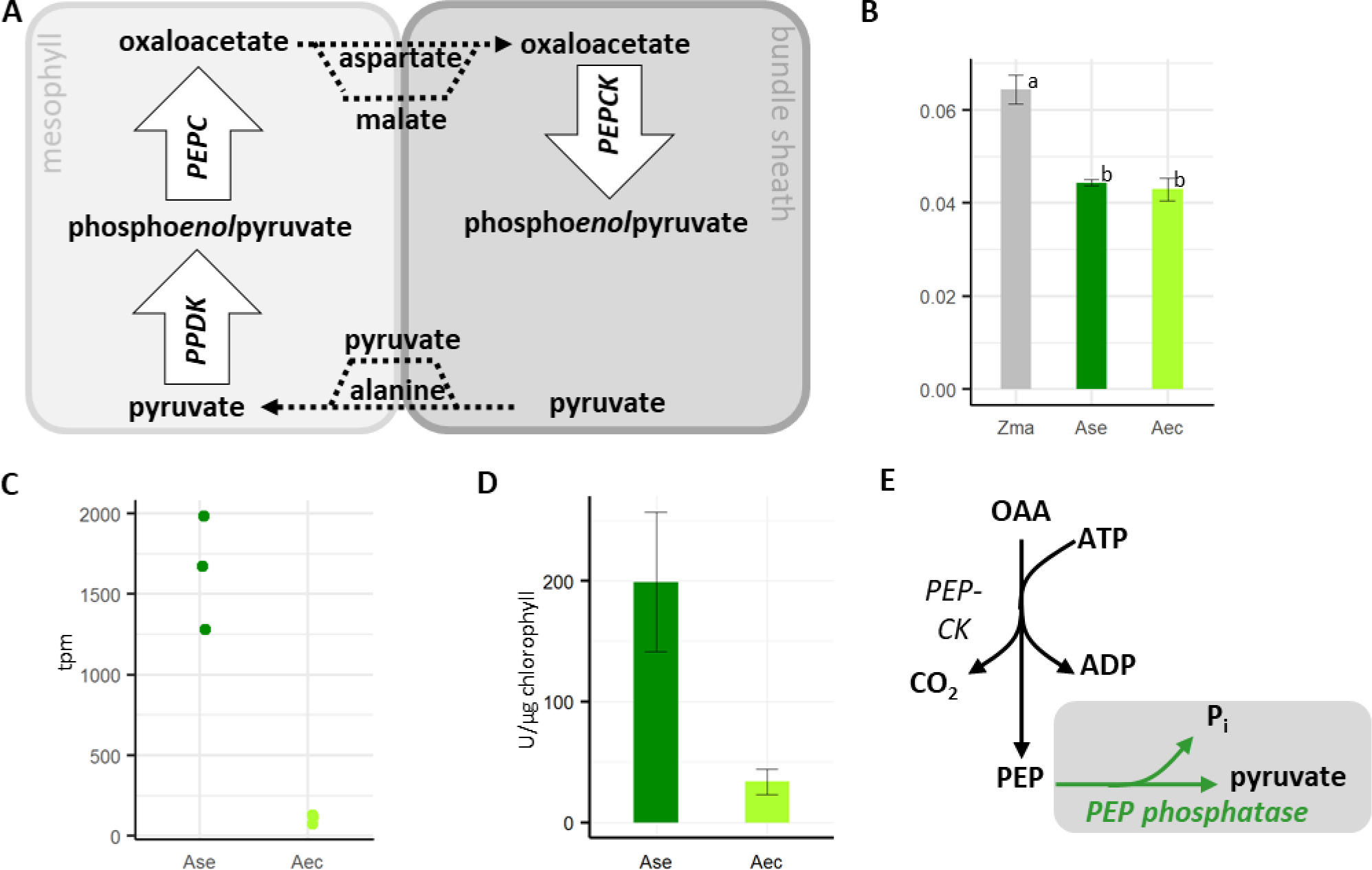
**The C_4_ cycle in PEP-CK dependent plants; (A)** schematic representation of a C_4_ cycle model without malic enzyme; PEPC PEP carboxylase, PEPCK PEP carboxykinase, PPDK pyruvate phosphate dikinase (**B**) quantum yield of the NADP-ME species *Z. mays*,the PEP-CKspeciesy *A.semialata* subsp. *semialata* (Ase), and a C_3_ species, *A. semialata* subsp. *eckloniana* (Aec);, error bars depict SEM, different letters denote significant differences (n=4, ANOVA followed by posthoc Tukey test) (**C**) expression of PEP phosphatase measured by cross species mapping onto *Setaria italica* transcriptome (gene ID Si021926m), expression is significantly different (edgeR, classic mode, Benjamini Hochberg corrected, n=3, p<0.01); Aec *Alloteropsis semialata subsp. eckloniana*, Ase *A. semialata subsp. semialata* (**D**) PEP phosphatase activity of leaf extract, activity is significantly different (t-test, error bars depict SEM, n=4); (**E**) PEP-CK in the cytoplasm is driven by irreversible removal of PEP in the PEP phosphatase reaction; OAA oxaloacetate, PEP phosphoenolpyruvate, driver is identified by a grey box

Comparative RNA-seq analyses between the C_4_*Alloteropsis semialata* subs. *semialata* and C_3_ subspecies *Alloteropsis semialata* subsp. *eckloniana* show highly abundant PEPC, PEP-CK, and PPDK transcripts (Supplemental Table 1). PPDK regenerates PEP from pyruvate in C_4_ species^8^, and implies the presence of pyruvate in the C_4_ cycle. Under the experimental conditions, however, malic enzyme abundance did not exceed 600 transcripts per million in the C_4_ individuals (Supplemental Figure 1, Supplemental Table 1) which confirmed earlier measurements of low malic enzyme activity^29,30^. Two candidate enzymatic reactions for closing the gap between PEP and pyruvate exist. Pyruvate kinase reversibly transfers the phosphate group from PEP to ATP thereby conserving the energy of the phosphate ester bond while forming pyruvate. Conversely, PEP phosphatase releases orthophosphate, resulting in a loss of chemical energy to entropy, which makes the reaction irreversible under physiological conditions.

The gap between pyruvate and PEP is apparently not closed by pyruvate kinase as none of these reach high expression levels (Supplemental Table 1, Supplemental Figure 2). *A. semialata* subs. *semialata* does not realize the considerable energy savings of a PEP-CK/pyruvate kinase decarboxylation model and has reduced quantum yield compared to *Z. mays* which relies on NADP-ME (Figure 3C). A transcript matching an experimentally validated PEP phosphatase^32^ shows increased abundance in *A. semialata* subsp. *semialata* (Figure 3C). Activity tests at cytosolic pH with whole leaf extracts detect high PEP phosphatase activity in *A. semialata* subs. *semialata* but not in its C_3_ sister *A. semialata* subsp. *eckloniana* (Figure 3D). Taken together, these results suggest that a PEP phosphatase closes the PEP-CK -based C_4_ cycle in *A. semialata* subs. *semialata* independent of malic enzymes. Unlike pyruvate kinase, which conserves the energy of the phosphate-ester bond, PEP phosphatase discards it, making the reaction irreversible. This additional difference in free reaction energy allows for high pyruvate concentrations required for the diffusion-driven transport back to the mesophyll cell while still maintaining an overall high decarboxylation rate. Thus, the irreversible dephosphorylation of PEP drives the PEP-CK based decarboxylation to high velocity (Figure 3E).

High velocity decarboxylation is critical to C_4_ photosynthesis since it is required to achieve the primary function of the cycle, the enrichment of CO_2_ at the site of Rubisco. We show that each of the decarboxylation enzymes is coupled to complementary biochemical reactions that accelerate the decarboxylation reactions. These drivers are not universal, but are instead specific to the particular enzyme, and provide explanations for well-documented^8^ yet hitherto unexplained C_4_ cycle features. For NADP-ME plants, chloroplastic redox poise requires a shift towards NADP. Previous attempts to explain the NADP-ME associated chloroplast dimorphism invoked selective pressure to limit O_2_ production near Rubisco^8^ but failed to explain why NAD-ME species never evolved the chloroplast dimorphism. Similarly, theoretical biochemistry always suggested malate as the most efficient transfer acid since it provided reduction power transfer^8^, making the predominance of aspartate as the transfer acid in NAD-ME species hard to explain^8^. Our driver models explain each observation as an enzyme-specific adaptation of the decarboxylation environment (Figure 2B, C). It also explains why the enzymes and transporters evolved so convergently in NADP-ME and NAD-ME types; the redox poise requirements in the organelles impose high constraints. Using PEP-CK as the sole or even dominant decarboxylation enzyme as observed in *Alloteropsissemilalata* (Figure 3, Supplemental Table 1, Supplemental Figure 1) was considered impossible up until now^8,26,28^. The identification of PEP-phosphatase not only closes a PEP-CK-based C_4_ cycle (Figure 3A) for the first time, but it also provides an explanation for the high decarboxylation drive. The essentially irreversible dephosphorylation of PEP to pyruvate (ΔG^0^< −61.5kJ/mol) provides a sink for PEP and drives high velocity PEP-CK based decarboxylation of oxaloacetate (Figure 3E).

By isolating the decarboxylation subsystems in models using only measured values as constraints rather than striving for models of the complete cycle constrained by all known features^3,26^, we identify the mechanistic roles of photosystem II depletion in NADP-ME-based C_4_ photosynthesis and transfer acid identity in NAD-ME-based C_4_ photosynthesis, and discover the enzyme that closes a rare predominantly PEP-CK -based C_4_ cycle in *A. semialata.* Testing for drivers of biochemical behavior in a mathematically isolated metabolic sub-system can enhance understanding of the phenotype beyond the modelled sub-system in unexpected ways.

Testing these conclusions will have to await the synthetic reconstruction of the pathway in a C_3_ plant chassis, in which the contributions of the adaptations for decarboxylation speed will become apparent.

## Acknowledgements

We acknowledge the European Union 7th Framework Program (EU 3to4 to APMW, CPO, and SB) and the Deutsche Forschungsgemeinschaft (EXC 1028 to OE) for financial support.

## Material and Methods

### Modeling

The model for NADP-ME was derived from^19^ according to^3^ using the following equations:

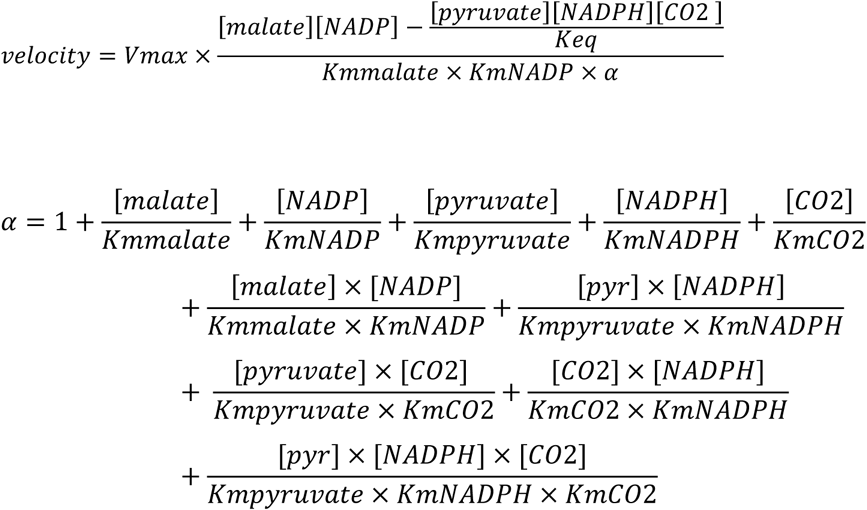

The model was parametrized from the literature: Whole bundle sheath cellular steady state concentrations of active pool malate and pyruvate were 5.2mM for malate and 0.26mM for pyruvate^16^. The CO_2_ concentration was estimated as 0.08mM from a partial pressure of 2400μbar in the bundle sheath in C4 plants^2^. The equilibration constant was calculated from ΔG°=14kJ/mol as K_eq_=0.0035M. The Km for malate was 0.39mM and the Km for NADP for the fully reduced activated enzyme at pH 8.0 was 0.0276mM^33^. Km for NADPH was 0.045mM, Km for pyruvate was 3mM and Km for CO_2_ was calculated at pH8.0 as 0.44mM from the Km HCO_3_^-^ of 22mM^34^. Total concentration of pyridine nucleotides was estimated as 0.14mM^35^. Photosynthetic rate (41μmol CO_2_ m^-2^s^-1^) and maximal NADP-ME activity (67μmol m^-2^s^-1^) were determined in mature leaves^17^. Running decarboxylation faster than Rubisco-catalyzed CO_2_-fixation is required to counteract leakage of CO_2_ from the bundle sheath cells. The overcycling rate required to achieve CO_2_ enrichment has been experimentally determined to be around 20%^2^. The code is available as an R script (Supplemental Data Rscript_model.R)

### Plant growth

*Alloteropsis* plants were grown under 23°C day/20°C night temperatures, 300 μmol m^-2^ s^-1^ photosynthetic photon flux density (PPFD), and ambient relative humidity and CO_2_ concentrations over 14h day/10h night cycles in John Innes No. 2 compost (John Innes Manufacturers Association, Reading, England), maintained at well-watered soil water conditions, and fertilized biweekly with Scotts Evergreen Lawn Food (The Scotts Company, Surrey, England).

### RNA-seq

Total RNA was extracted from climate chamber grown plants at the midpoint of the light cycle using the RNeasy Plant Mini Kit (Qiagen, Hilden, Germany). Libraries were prepared (TruSeq RNA sample preparation kit, Illumina, San Diego) after rapid DNase digest and sequenced on a HiSeq2000 sequencer (Biologisch-Medizinisches Forschungszentrum of the Heinrich Heine University Düsseldorf, Germany) as paired-end reads in three biological replicates. The reads were aligned to the *Setaria italica* transcriptome (phytozome V11) with Blat^36^ according to^11^ and transformed to transcripts per million for plotting. Significantly differently expressed references were determined with edgeR on raw read counts^37^ and FDR corrected^38^.

### Enzyme measurements

PEP phosphatase activity was measured in biological quadruplicates. 20-50 mg of powdered plant material were extracted in extraction buffer (50mM Hepes, pH7.5, 5mM MgCl_2_, 2mM MnCl_2_, 5mM DTT, 2mM EDTA, 10% glycerin, and 0.1% tritonX100) by brief vortexing. The plant material was pelleted and the supernatant used for analysis. PEP phosphatase was assayed by following NADH oxidation at 340nm in assay buffer (100mM Hepes pH 6.9, 50mM KCl, 10mM MgCl_2_, 0.2mM NADH, 2mM DTT, 0.2mg/ml BSA, 2U lactate dehydrogenase) after starting the reaction with 2mM PEP^39^. Pyruvate kinase activity was determined by including ADP in the reaction buffer and was not detectable against the background of PEP phosphatase activity. The initial slope, averaged over two technical replicates, was used to calculate the consumption of NADH per chlorophyll content^40^.

### Quantum yield measurements

*Z.mays, A. semialata* subsp. *semialata,* and *A. semialata* subsp. *ecklonia* were grown under greenhouse conditions controlled to 25/20°C day/night temperatures over a 12 h photoperiod supplemented with constant artificial light to achieve a minimum light intensity of 200 μmol m^-2^ s^-1^, at ambient relative humidity and CO_2_. On four replicate plants per species, young fully expanded leaves were measured using an open gas exchange system with an infra-red gas analyzer (LI-6400XT and LI-6400-40, respectively; LICOR, Lincoln, Nebraska, USA). Net carbon assimilation was measured at a gradient of low light intensities (100, 90, 80, 70, 60, 50, 40, 30, 20, 10, 0 μmol m^-2^ s^-1^), at 27°C leaf temperature, 250 μmol s^-1^ flow rate, sample cell CO_2_ concentration (C_a_) of 400 μmol CO_2_ mol^-1^, and approximately 1.2 kP leaf temperature vapour pressure deficit, after sufficient acclimation to chamber conditions to achieve steady state CO_2_ and H_2_O fluxes. Leaf absorbance was estimated from measurements of leaf transmittance per LI-COR technical note 128 protocol and averaged over four replicate plants per species. Quantum yield of CO_2_ was calculated as the rate of assimilated CO_2_ per number of absorbed photons.

### Data availability

RNA-seq data generated during this study is available at ENA as SRXXXXXX-XX. Processed RNA-seq data is available as a supplemental table including results of statistical tests.

**Author contributions**: AB conceived the driver concept, implemented the models, did the RNA-seq experiments, analyzed the results and wrote the paper, US analyzed RNA-seq results and discussed the models, MRL measured quantum yield, SF measured PEP phosphatase activity, OE and GS assisted in model implementation, PAC and MRL assisted in RNA-seq and activity measurements, SB and JMD processed RNA-seq data, APMW and CPO conceived the RNA-seq approach, UG discussed the models, assisted in implementation and co-wrote the paper. All authors commented on the manuscript and approved the final submission.

